# Effects of clozapine-N-oxide and compound 21 on sleep in laboratory mice

**DOI:** 10.1101/2022.02.01.478652

**Authors:** Janine Traut, Jose Prius Mengual, Elise J. Meijer, Laura E. McKillop, Hannah Alfonsa, Anna Hoerder-Suabedissen, Seoho Michael Song, Zoltán Molnár, Colin J. Akerman, Vladyslav V. Vyazovskiy, Lukas B. Krone

## Abstract

Designer Receptors Exclusively Activated by Designer Drugs (DREADDs) are chemogenetic tools for remote control of targeted cell populations using chemical actuators that bind to modified receptors. Despite the popularity of DREADDs in neuroscience and sleep research, potential effects of the DREADD actuator clozapine-N-oxide (CNO) on sleep have never been systematically tested. Here we show that intraperitoneal injections of commonly used CNO doses (1, 5, and 10 mg/kg) alter sleep in wild-type male laboratory mice. Using electroencephalography (EEG) and electromyography (EMG) to analyse sleep, we found a dose-dependent suppression of rapid eye-movement (REM) sleep, changes in EEG spectral power during non-REM (NREM) sleep, and altered sleep architecture in a pattern previously reported for clozapine. Effects of CNO on sleep could arise from back-metabolism to clozapine or binding to endogenous neurotransmitter receptors. Interestingly, we found that the novel DREADD actuator, compound 21 (C21, 3 mg/kg), similarly modulates sleep despite a lack of back-metabolism to clozapine. Our results demonstrate that both CNO and C21 can modulate sleep of mice not expressing DREADD receptors. This implies that back-metabolism to clozapine is not the sole mechanism underlying side effects of chemogenetic actuators. Therefore, any chemogenetic experiment should include a DREADD-free control group injected with the same CNO, C21 or newly developed actuator. We suggest that electrophysiological sleep assessment could serve as a sensitive tool to test the biological inertness of novel chemogenetic actuators.

## Introduction

Chemogenetics is an important and widely used experimental approach in sleep research (Weber and Dan 2016; Varin and Bonnavion 2019) and could serve as a novel therapeutic strategy in sleep medicine (Venner et al. 2019). Designer Receptors Exclusively Activated by Designer Drugs (DREADDs) enable non-invasive and cell-type specific remote control of neuronal activity in freely moving animals on a time scale of minutes to hours (Roth 2016). In the context of sleep research, these characteristics make DREADD technology a powerful tool with which to probe the contribution of selected neuronal populations in controlling vigilance states (Hayashi et al. 2015; Varin, Luppi, and Fort 2018; Yu et al. 2019; Mondino et al. 2021), sleep-state specific network oscillations (Funk et al. 2017; Vaidyanathan et al. 2021), as well as sleep-related physiology (Harding et al. 2018; Fleury Curado et al. 2018) and behaviour (Eban-Rothschild et al. 2016; Tossell et al. 2020).

Typically, intraperitoneal injections of clozapine-N-oxide (CNO) are used to activate excitatory hM3Dq or inhibitory hM4Di DREADDs (Campbell and Marchant 2018). Early work suggested that CNO is pharmacologically inert (Armbruster et al. 2007) and not back metabolized to its parent drug clozapine in mice (Guettier et al. 2009). However, more recent studies demonstrated relevant conversion of CNO to pharmacologically active metabolites including clozapine (Gomez et al. 2017; Manvich et al. 2018; Jendryka et al. 2019). Clozapine is an atypical antipsychotic drug used in the treatment of schizophrenia with a high affinity for dopamine D2 and serotonin 5-HT2A coupled with a broad binding profile to cholinergic, adrenergic, histaminergic and serotonergic receptors (Wenthur and Lindsley 2013), which may account for its high efficacy compared to other antipsychotics (Kane et al. 1988). In addition, CNO itself was found to present off-target binding at a broad range of neurotransmitter receptors (Gomez et al. 2017; Jendryka et al. 2019) and to elicit behavioural effects (Gomez et al. 2017; Manvich et al. 2018; MacLaren et al. 2016) at doses commonly used for DREADD experiments.

It is widely thought that CNO does not affect sleep (Harding et al. 2018; Mondino et al. 2021; Naganuma et al. 2018; Erickson et al. 2019) and many chemogenetic sleep studies include convincing control experiments with DREADD-free animals that demonstrate the absence of relevant effects of the chosen CNO preparations and doses on the assessed sleep parameters (Erickson et al. 2019; Mondino et al. 2021; Takata et al. 2018; Yu et al. 2019; Anaclet et al. 2015; Venner et al. 2016). However, a comprehensive assessment of putative dose-dependent effects of CNO on sleep in wild-type mice has never been systematically conducted, although a recent study reported that high CNO doses affected sleep in DREADD-free control animals (Varin, Luppi, and Fort 2018). This is an important omission, given that clozapine is known to modulate sleep in humans (Hinze-Selch et al. 1997; Monti, Torterolo, and Pandi Perumal 2017; Riemann and Nissen 2012) and laboratory rodents (Spierings, Dzoljic, and Godschalk 1977; Sorge, Pollmächer, and Lancel 2004; Grønli et al. 2016; Coward et al. 1989). Many of the endogenous neurotransmitter receptors, which are drug targets of clozapine (Wenthur and Lindsley 2013) and to which CNO presents off-target binding affinity (Jendryka et al. 2019; Gomez et al. 2017), are also involved in the regulation of arousal and sleep (Saper and Fuller 2017).

Initially intended as a control experiment, we here tested whether commonly used doses of CNO (1, 5, and 10 mg/kg) and the novel DREADD agonist compound 21 (C21), which does not convert to clozapine (Thompson et al. 2018) but has an off-target binding profile similar to CNO (Jendryka et al. 2019), affect sleep in wild-type C57BL/6J mice under laboratory conditions. We find dose-dependent clozapine-like effects of CNO on the duration of rapid eye movement (REM) sleep, sleep architecture parameters, and frontal EEG power spectra of non-rapid eye movement (NREM) sleep. In addition, we observed a similar pattern of sleep modulation after injections of a 3 mg/kg dose of C21 resulting in effect sizes comparable to those of the 5 mg/kg CNO condition.

## Results

### Clozapine-N-oxide suppresses REM sleep

We first assessed the time spent in wakefulness, non-rapid eye movement (NREM) and rapid eye-movement (REM) sleep in hourly intervals over the first six hours of the light period following injections of CNO or saline at light onset (Figure 1a). There was no significant main effect of the treatment condition on the proportion of time spent awake or in NREM and no interaction effect between treatment condition and time point. However, we observed a significant main effect of the treatment condition on the percentage of REM sleep (*F*_(2, 30)_ = 5.601, *p* = 0.009), but no interaction between the time point and the treatment condition (Figure 1a).

**Figure 1:**
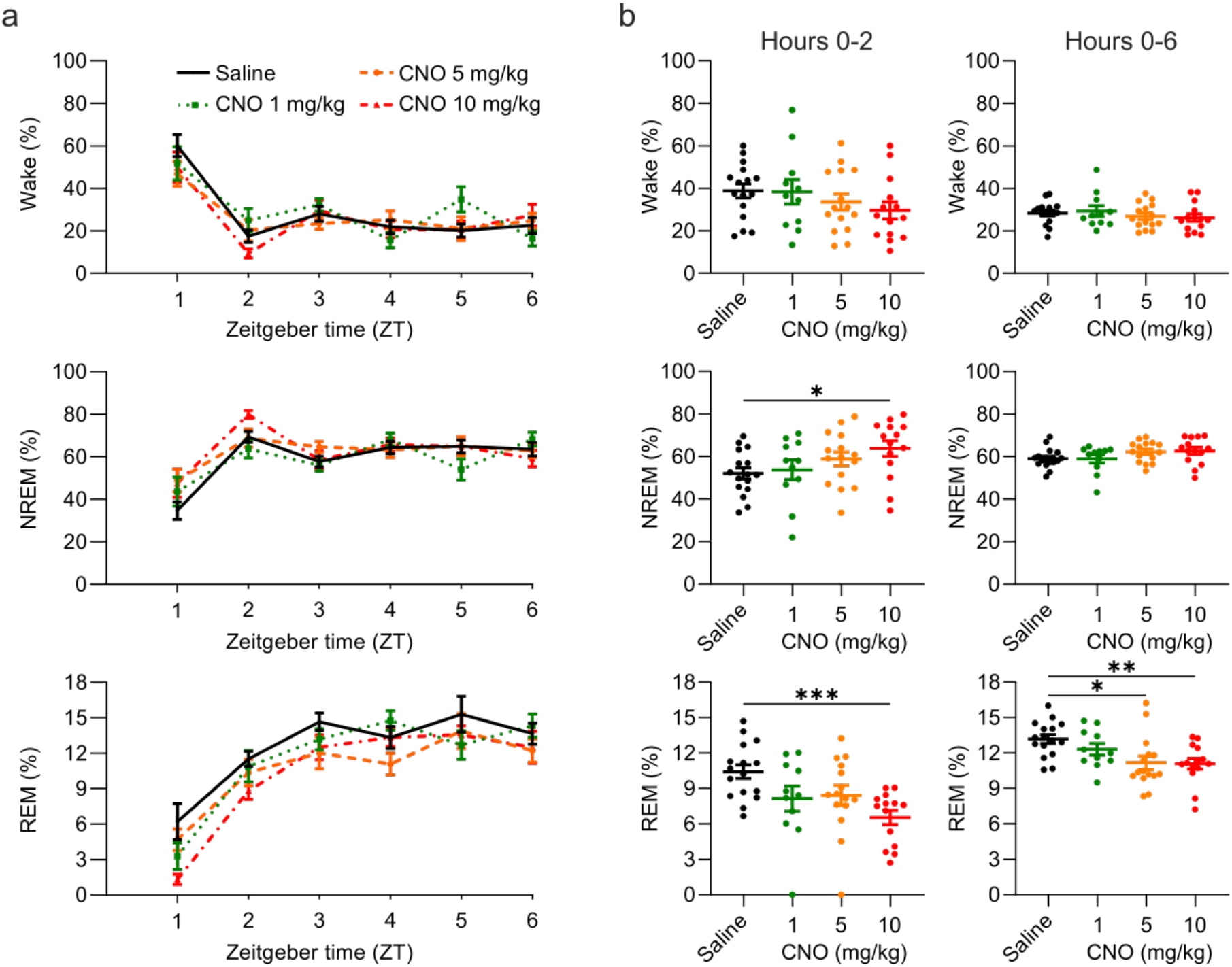
Suppression of REM sleep following CNO injection. a) Time course of wakefulness, NREM, and REM sleep in the six hours following subcutaneous injections of CNO or saline at light onset (ZT 0). b) Percentage of time spent in the three vigilance states during the first two hours (left column) and over the entire six-hour observation period after saline and CNO injections. Note that there is no main effect of the treatment condition on NREM sleep in the acute time window (*F*_(2,24)_ = 3.054, *p* = 0.068). Therefore, the significant post-hoc contrast for the 10 mg/kg condition has to be interpreted with caution. Percentage of REM sleep expressed as proportion of total sleep time. n=16 for saline, n=11 for 1 mg/kg, n=15 for 5 mg/kg, n=14 for 10 mg/kg for vigilance state analysis. Asterisks indicate post-hoc contrasts with significant differences (*P < 0.05, **P < 0.01, ***P < 0.001). CNO: clozapine-N-oxide. EEG: electroencephalogram. NREM: non-rapid eye movement sleep. REM: rapid eye movement sleep. ZT: Zeitgeber time.

CNO concentrations in blood plasma, cerebrospinal fluid (CSF) and brain tissue of mice peak within the first 15-30 minutes after both intraperitoneal and subcutaneous injections (Jendryka et al. 2019; Manvich et al. 2018), and behavioural side effects are typically tested within the first two hours following drug administration (MacLaren et al. 2016; Manvich et al. 2018; Gomez et al. 2017). Yet, CNO effects might persist for much longer. To investigate this possibility more closely, we next analysed the effect of CNO on vigilance states in an acute (0-2 hours) and prolonged (0-6 hours) time window (Figure 1b). There was a significant effect of CNO treatment on REM sleep both in the acute (*F*_(2, 22)_ =8.951, *p* = 0.002) and prolonged time window (*F*_(2, 23)_ = 7.525, *p* = 0.004). Effect size calculations for the post-hoc comparisons between the individual CNO conditions and the saline condition indicated medium to large effects of CNO on REM sleep (Supplementary Table 1).

### Clozapine-N-oxide alters sleep architecture

Physiological sleep in mammals is typically entered through NREM sleep and characterised by the alternation between NREM and REM sleep episodes. The average timing, duration, and frequency of NREM and REM episodes in mice vary slightly depending on the genetic background but are kept within tight limits for individual strains (Franken, Malafosse, and Tafti 1999; Huber, Deboer, and Tobler 2000). The effects of psychotropic drugs on sleep are often most prominent in sleep architecture parameters (Riemann and Nissen 2012). For example, clozapine evokes characteristic changes in sleep architecture in humans (Monti, Torterolo, and Pandi Perumal 2017), rats (Sorge, Pollmächer, and Lancel 2004; Spierings, Dzoljic, and Godschalk 1977), and mice (Grønli et al. 2016). Hence, the analysis of sleep architectural parameters is of paramount importance in assessing whether a pharmacological compound modulates sleep.

Characteristics of both NREM and REM episodes were altered by CNO injections (Figure 2a). CNO injections were followed by longer (main effect NREM episode duration: *F*_(2, 30)_ = 11.64, *p* < 0.001) but fewer (main effect NREM episode number: *F* _(2, 31)_ = 7.796, *p* = 0.001) NREM sleep episodes. These effects appeared to be dose-dependent, with higher doses eliciting stronger effects (Figure 2b). The latency between the injection and the onset of NREM sleep was also significantly affected by CNO injections (*F*_(2,26)_ = 3.380, *p* = 0.046, Figure 2b). However, the reduction of the NREM sleep latency was only statistically significant in the 1 mg/kg condition. REM sleep episodes were significantly shorter following CNO injections (*F*_(2, 40)_ = 4.509, *p* = 0.013) and the latency between sleep onset and the first transition to REM sleep was significantly changed (*F*_(2, 28)_ = 6.985, *p* = 0.002) with delayed REM onset in the 1 mg/kg and 10 mg/kg condition (Figure 2b). In contrast to the effects on sleep states, neither the duration nor the number of wake episodes was modulated by CNO injections (Supplementary Table 2). In summary, we observed longer but fewer NREM episodes following CNO injections in a dose-dependent fashion, and a similar but less pronounced change in the duration and frequency of REM episodes. Furthermore, CNO injections accelerated sleep onset, but delayed the transition to REM sleep, particularly at low doses.

**Figure 2:**
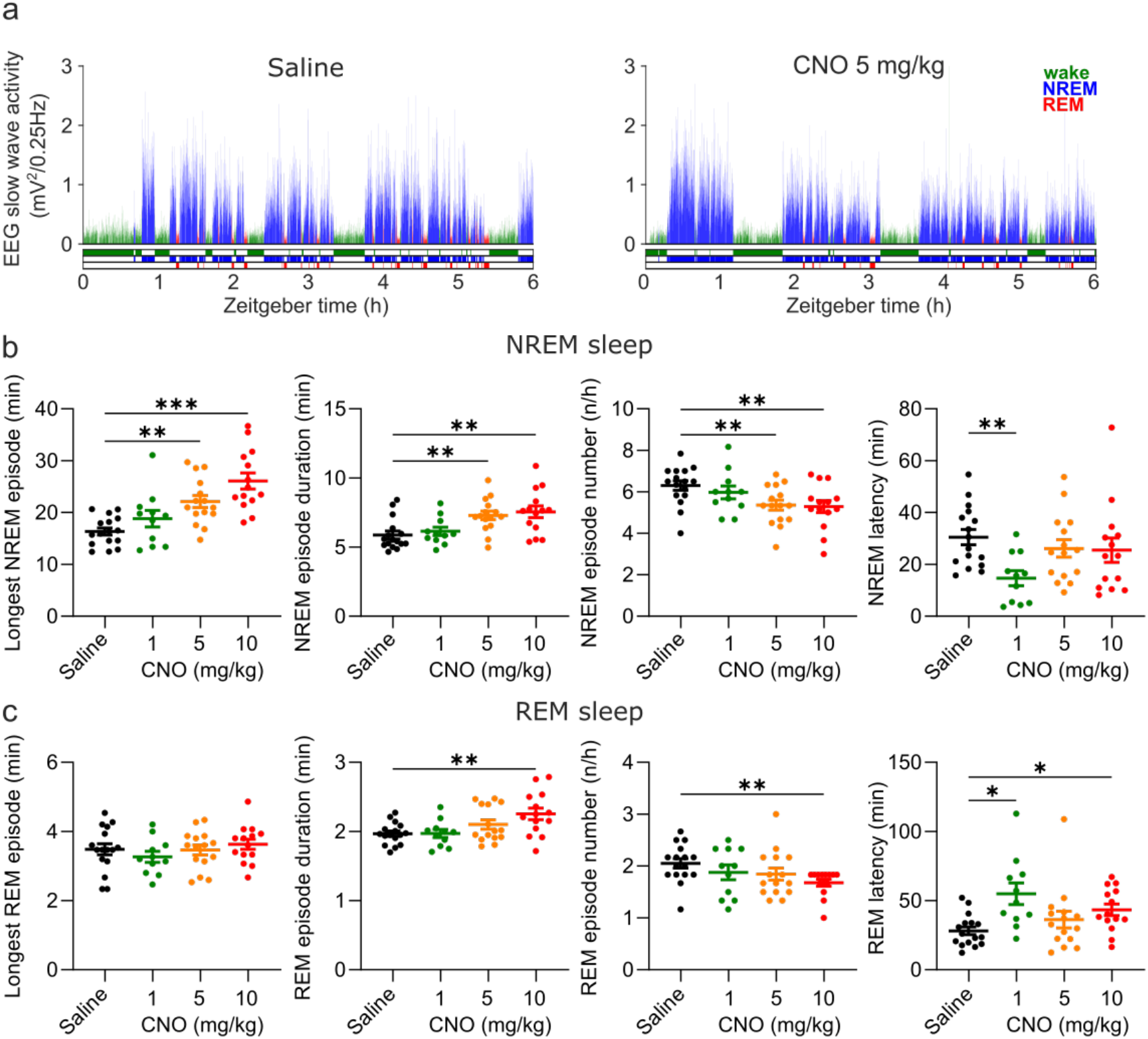
Altered sleep architecture following CNO injections. a) Representative hypnograms and EEG slow wave activity (0.5-4.0 Hz, 4-s epochs) from one individual mouse after injection of saline (left panel) and 5 mg/kg CNO (right panel). Note the reduced latency to NREM sleep, the suppression of REM sleep, and the increased duration of individual NREM sleep episodes. b) NREM sleep architecture following saline and CNO injections. c) REM sleep architecture following saline and CNO injections. Note that there is no main effect of CNO treatment on the number of REM sleep episodes (*F*_(3,33)_ = 2.680, *p* = 0.069). Therefore, the significant post-hoc contrast for the 10 mg/kg condition has to be interpreted with caution. n=16 for saline, n=11 for 1 mg/kg, n=15 for 5 mg/kg, n=14 for 10 mg/kg for vigilance state analysis in panels b and c. Asterisks indicate post-hoc contrasts with significant differences (*P < 0.05, **P < 0.01, ***P < 0.001). CNO: clozapine-N-oxide. EEG: electroencephalogram. NREM: non-rapid eye movement sleep. REM: rapid eye movement sleep.

### Clozapine-N-oxide affects the NREM spectrogram and sleep consolidation

In addition to sleep time and architecture, EEG spectra are typically analysed in sleep studies. We therefore assessed whether CNO affects EEG spectral power. The focus of the EEG spectral analysis was on the comparison between the medium dose (5 mg/kg) of CNO and saline during NREM sleep. We observed that CNO injections were followed by a small but significant increase in spectral power in the range between 0.5 and 1.25 Hz and suppression of spectral power in nearly all frequency bins between 6 and 30 Hz during NREM sleep over the first two hours (Figure 3a and Supplementary Figure 1). While the increase in slow frequency bins during NREM sleep appeared to be temporary, the suppression of power above 6 Hz was significant for the entire six-hour observation period. Spectral analysis of wakefulness and REM sleep did not reveal any systematic effects of CNO (Supplementary Figure 1). A comparison of all three CNO conditions with saline injection indicated similar effects of CNO on the NREM sleep spectrogram in the high dose (10 mg/kg CNO), but not in the low dose (1 mg/kg CNO) condition (Figure 1).

**Figure 3:**
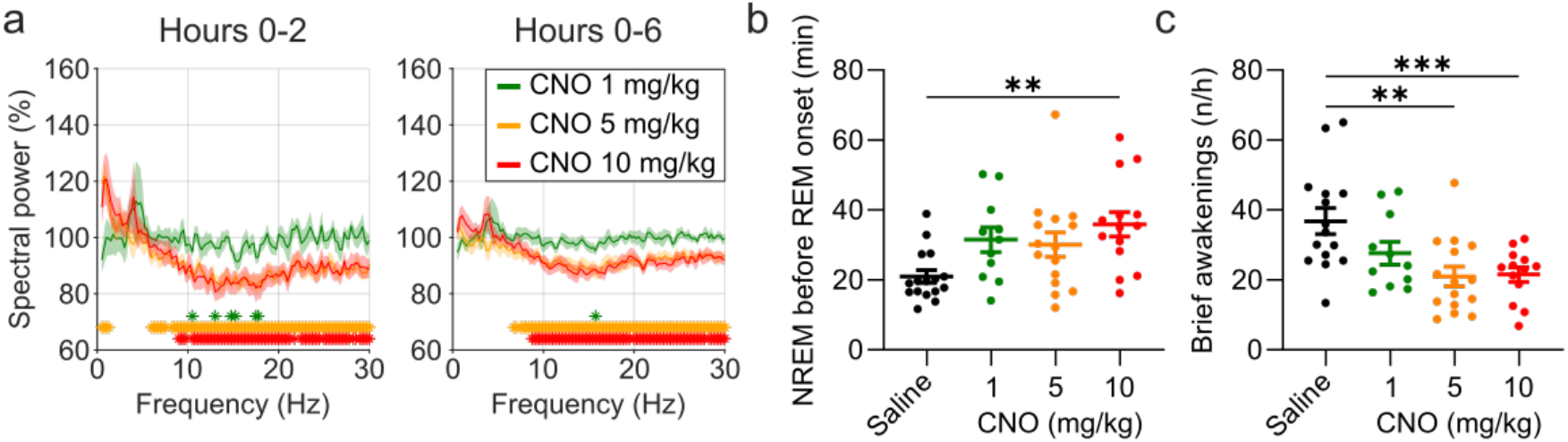
EEG spectral changes and increased sleep continuity following CNO injections. a) Frontal EEG spectra during NREM sleep relative to saline injections for the acute (first 2 hours) and prolonged (6 hours) observation period. b) Cumulative amount of NREM sleep before the first REM sleep episode. c) Frequency of brief awakenings (4-16 sec) per hour of sleep for the first 2 hours after injections. n=10 for saline, n=6 for 1 mg/kg, n=10 for 5 mg/kg, n=8 for 10 mg/kg for spectral analysis. n=16 for saline, n=11 for 1 mg/kg, n=15 for 5 mg/kg, n=14 for 10 mg/kg for vigilance state analysis. n=15 for saline, n=11 for 1 mg/kg, n=15 for 5 mg/kg, n=13 for 10 mg/kg for analysis of brief awakenings. Asterisks in panels b and c indicate post-hoc contrasts with significant differences (*P < 0.05, **P < 0.01, ***P < 0.001). Asterisks in panel a indicate frequency bins with significant differences in post-hoc comparison (P < 0.05). Data in a are presented as the mean ± s.e.m. (shaded areas). CNO: clozapine-N-oxide. EEG: electroencephalogram. NREM: non-rapid eye movement sleep.

Elevated spectral power in slow frequencies, longer but fewer NREM episodes, and less REM sleep are typically observed during the initial recovery sleep following sleep deprivation, when sleep is more consolidated (Huber, Deboer, and Tobler 2000). To explore whether the stability of sleep is affected by CNO, we assessed the cumulative amount of NREM sleep before the first REM episode (1 min or longer of REM sleep), as well as the frequency of brief awakenings (4-16 second intrusions of wake-like EMG and EEG during sleep), which is a behavioural marker of sleep continuity (Franken et al. 1991). Both measures were significantly changed after CNO injection (NREM before REM: *F*_(3, 33)_ = 5.071, *p* = 0.007; brief awakenings: *F*_(2, 24)_ = 10.38, *p* < 0.001). Effect size calculations indicated a medium to strong increase in NREM sleep before REM onset and reduction in brief awakenings for all CNO doses (Supplementary Table 2).

### Compound 21 has sleep-modulatory effects similar to clozapine-N-oxide

Based on recent reports indicating that back-metabolism of CNO to clozapine causes behavioural effects in rodents (Ilg et al. 2018; Gomez et al. 2017; Manvich et al. 2018; MacLaren et al. 2016) and might affect sleep (Varin, Luppi, and Fort 2018), we postulated that the effects of CNO injections on sleep could be avoided by using alternative DREADD ligands. However, another possibility, is that it is the off-target binding of CNO to endogenous receptors that causes or contributes to the change in sleep patterns. In this scenario, the effects of CNO on sleep would be, at least in part, mediated by direct action of the DREADD actuator at neurotransmitter receptors, which are involved in the regulation of sleep and could not be overcome by minimising the conversion to clozapine. To discriminate between these two possibilities, we investigated whether the next generation DREADD actuator C21 (Chen et al. 2015) has sleep-modulatory effects similar to those observed after CNO injections. C21 does not back-convert to clozapine *in vivo* (Thompson et al. 2018) but has an almost identical profile of binding affinities to endogenous neurotransmitter receptors as CNO (Jendryka et al. 2019).

Intraperitoneal injections of C21 at a dose of 3 mg/kg modulated sleep compared to saline (Figure 4 and Figure 5). REM sleep was significantly reduced over the six-hour observation period (*t*_(6)_ = 3.234, *p* = 0.009, Cohen’s d = -1.22). The numerical reduction of REM sleep in the acute, two-hour, time window following C21 injections did not reach statistical significance in this small sample of seven mice (*t*_(6)_ = 1.829, *p* = 0.059, Cohen’s d = -0.70; Figure 4b), but had an effect size comparable to that of the statistically significant REM sleep reduction in the same time window after 5 mg/kg CNO injections.

**Figure 4:**
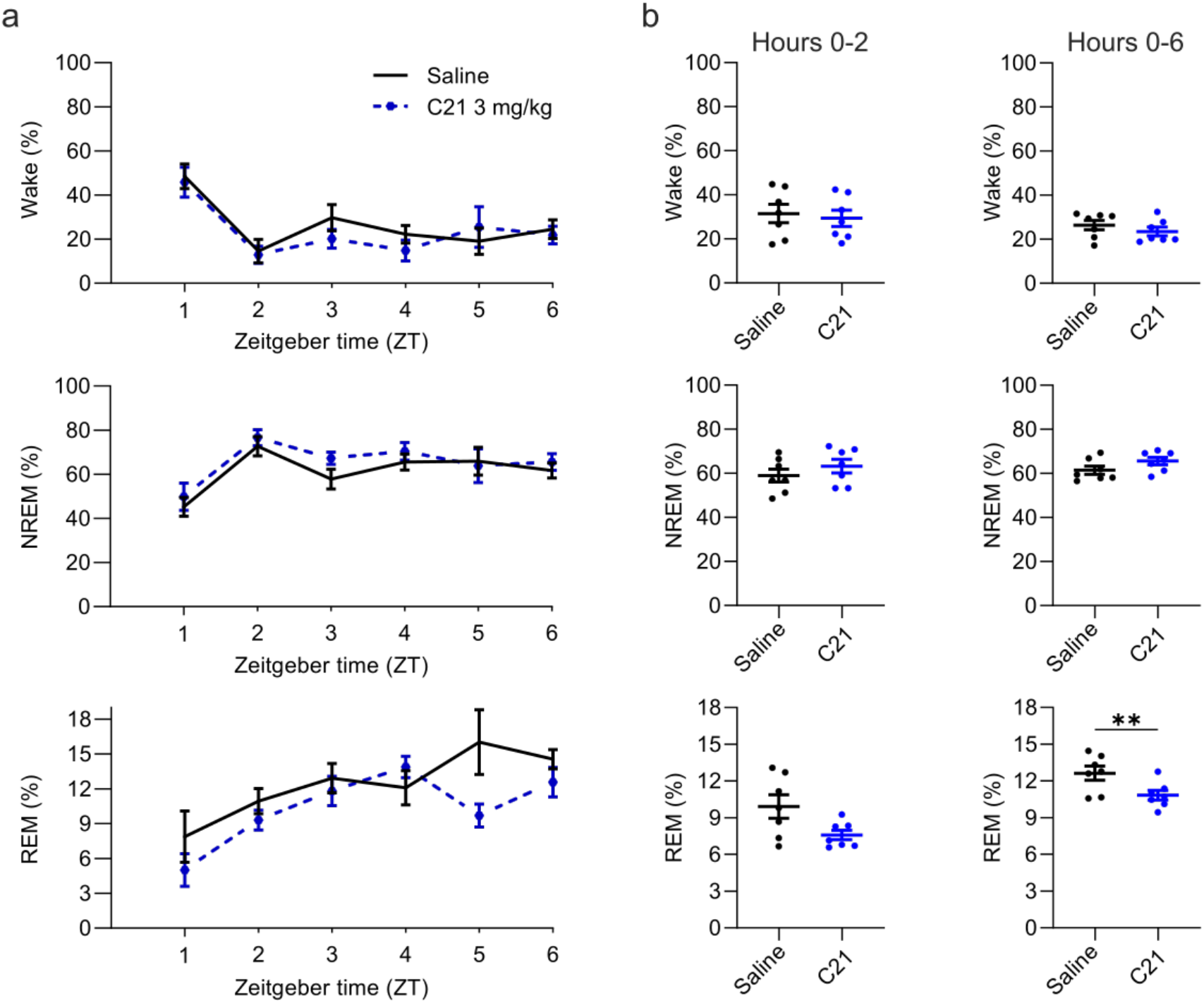
Suppression of REM sleep following C21 injections. a) Time course of wakefulness, NREM, and REM sleep in the six hours following injection of C21 or saline at light onset (ZT 0). b) Percentage of time spent in the three vigilance states during the first two hours (left column) and over the entire six-hour observation period after saline and C21 injections. Percentage of REM sleep expressed as proportion of total sleep time. n=7. Asterisks indicate post-hoc contrasts with significant differences (*P < 0.05, **P < 0.01, ***P < 0.001). C21: compound 21. EEG: electroencephalogram. NREM: non-rapid eye movement sleep. REM: rapid eye movement sleep. ZT: Zeitgeber time.

**Figure 5:**
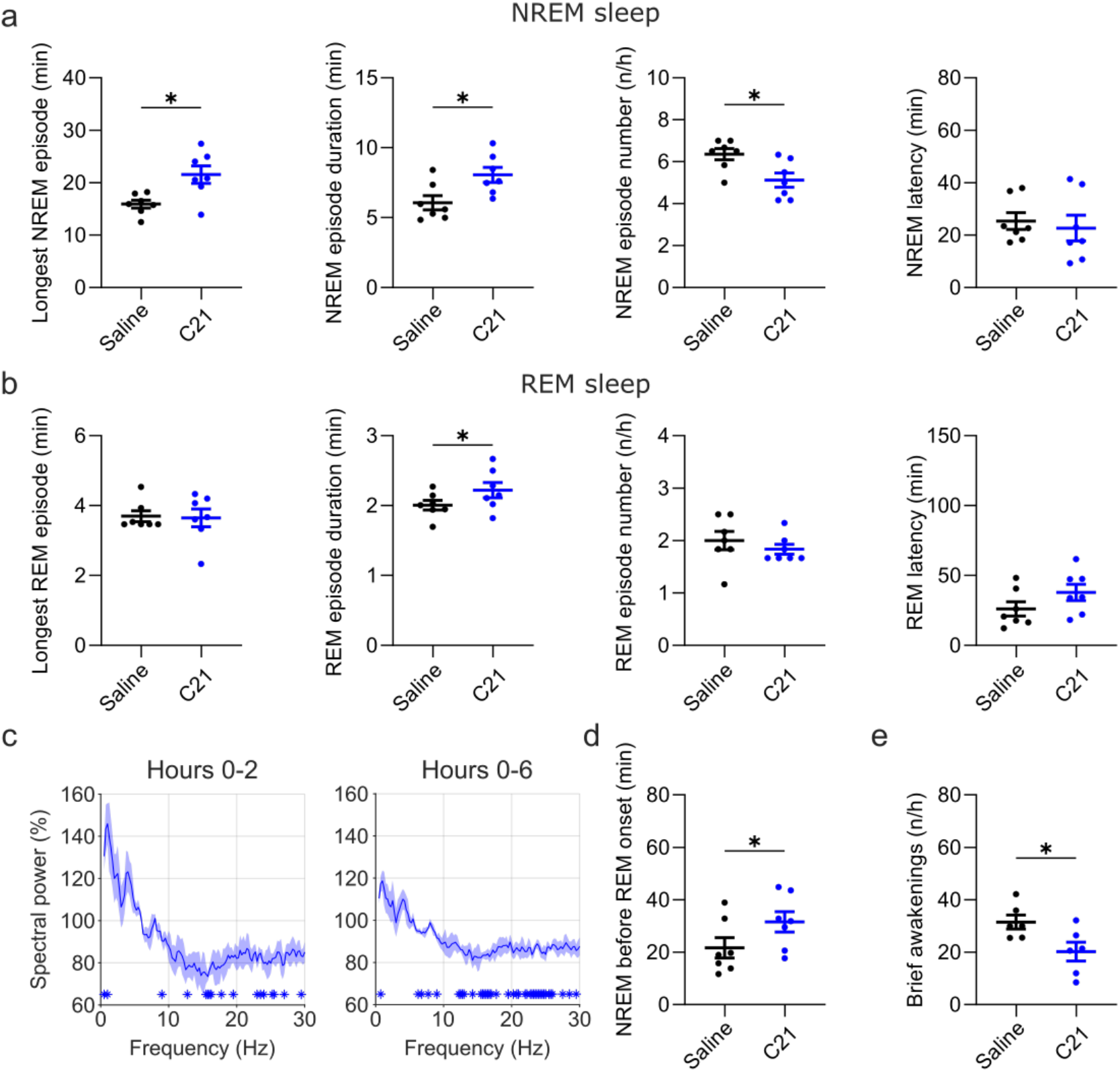
Changes in sleep architecture, NREM sleep spectra, and sleep continuity measures following C21 injections. a) NREM sleep architecture. b) REM sleep architecture. c) Frontal EEG spectra during NREM sleep relative to saline injections for the acute (2 hour) and full (6 hour) observation period following C21 injections. d) Cumulative amount of NREM sleep before the first REM sleep episode. e) Frequency of brief awakenings (4-16 sec) per hour of sleep for the first 2 hours after injections. Number of animals n=7 mice for vigilance state analysis in panels a, b and d. For spectral analysis in panel c: n=3. For analysis of brief awakenings in panel e: n=6 mice. Asterisks in panels a, b, d and e indicate post-hoc contrasts with significant differences (*P < 0.05, **P < 0.01, ***P < 0.001). Asterisks in panel c indicate frequency bins with significant differences in post-hoc comparison (P < 0.05). Data in c are presented as the mean ± s.e.m. (shaded areas). C21: compound 21. EEG: electroencephalogram. NREM: non-rapid eye movement sleep. REM: rapid eye movement sleep.

In addition to the suppression of REM sleep, C21 elicited significant changes in sleep architecture (Figure 5a,b). The maximum and average duration of NREM sleep episodes was increased (maximum duration: *t*_(6)_ = 2.551, *p* = 0.022, Cohen’s d = 0.96; average duration: *t*_(6)_ = 2.462, *p* = 0.025, Cohen’s d = 0.93) while the number of NREM sleep episodes in the first six hours following C21 injection was reduced (*t*_(6)_ = 2.809, *p* = 0.015, Cohen’s d = -1.06; Figure 5a). The average duration of REM episodes was also increased (*t*_(6)_ = 2.229, *p* = 0.034, Cohen’s d = 0.84; Figure 5b). For all NREM and REM architecture parameters that were significantly changed after CNO injections, effect sizes of 3 mg/kg C21 were similar to that of 5 mg/kg CNO compared to saline, even if the post-hoc comparisons did not reach significance in this small sample (*Supplementary Table 3*). NREM spectra showed an increase in the very slow frequency bins and a suppression of frequencies above 6 Hz (Figure 5c) – a pattern which resembled that of the medium and high dose CNO injections (Figure 3a). Markers of sleep consolidation were also significantly altered as a result of C21 injections. The amount of NREM sleep before the first occurrence of REM sleep was increased (t_(6)_ = 2.252, *p* = 0.033, Cohen’s d = 0.85) and brief awakenings were reduced (*t*_(5)_ = 2.164, *p* = 0.041, Cohen’s d = -0.52; Figure 5d,e).

## Discussion

### CNO and C21 injections have clozapine-like effects on sleep

Our study demonstrates that the chemogenetic actuators CNO and C21 can modulate sleep in wild-type laboratory mice which do not express DREADD receptors. Both substances led to a suppression of REM sleep and affected sleep architecture and NREM sleep spectra in a pattern consistent with more consolidated sleep. The sleep changes following CNO and C21 injections in wild-type mice bore striking similarities with those previously reported for clozapine in rats (Sorge, Pollmächer, and Lancel 2004; Spierings, Dzoljic, and Godschalk 1977), and humans (Hinze-Selch et al. 1997). In particular, the initial suppression of REM sleep and the occurrence of longer but fewer NREM episodes were the most consistent dose-dependent effects of clozapine on sleep in male Wistar rats, leading the authors to conclude that clozapine has sedative effects, suppresses REM initiation and increases sleep maintenance in rats (Sorge, Pollmächer, and Lancel 2004). Interestingly, in rats low doses of clozapine (2.5 mg/kg) had immediate sleep-promoting effects while high doses of clozapine (7.5 mg/kg) initially promoted wakefulness before a sustained increase NREM sleep after the first two hours (Sorge, Pollmächer, and Lancel 2004). This might explain why in our study NREM and REM sleep latency were most strongly affected by the lowest dose (1 mg/kg) of CNO. Our findings of sleep modulatory effects of CNO are in line with previous reports of behavioural side effects of CNO resulting from back-metabolism to clozapine (Gomez et al. 2017; Manvich et al. 2018). However, the fact that we observed similar changes in sleep after injections with C21, a DREADD actuator that does not convert to clozapine (Thompson et al. 2018), suggests that *in vivo* metabolism to clozapine conversion is not the sole mechanism through which DREADD actuators can elicit unwanted effects.

### Possible mechanisms underlying sleep-modulatory effects of CNO and C21

In our view, the most parsimonious explanation for the observed sleep-modulatory effects of CNO and C21 is off-target binding at endogenous neurotransmitter receptors. It has been shown that CNO is a competitive inhibitor of several neurotransmitter receptors, including histaminergic H1, serotoninergic 5-HT1A, 5-HT1B, 5-HT2A, 5-HT2B, muscarinic M1, M2, M3, M4, adrenergic α1A and α2A, and dopaminergic D1 and D2 receptors (Gomez et al. 2017; Jendryka et al. 2019). C21 has an off-target binding profile similar to CNO (Jendryka et al. 2019). In addition, a recent study reported increased firing rates of nigral dopaminergic neurons in wild-type rats, indicating that C21 can elicit direct neuromodulatory effects in rodents (Goutaudier et al. 2020). Among several endogenous receptors relevant for sleep regulation, CNO and C21 induce a strong competitive inhibition at histamine H1-receptors (Jendryka et al. 2019). Tested against a panel of g-protein coupled receptors (GPRCs), C21 had a greater affinity for histamine H1-receptors than for muscarinic DREADDs (Thompson et al. 2018). H1-receptor knockout and pharmacological antagonism of the H1 receptor in mice both result in a reduced number of brief awakenings, fewer but longer NREM sleep episodes, and a reduced latency to NREM sleep (Huang et al. 2006). In addition, antihistamines which induce drowsiness, are known to strongly suppress REM sleep, but can be acutely NREM-promoting at low doses and wake-promoting at high doses, respectively (Ikeda-Sagara et al. 2012). Therefore, we speculate that the shared sleep-modulatory effects of CNO and C21 might be due to direct antihistaminergic action. However, a contribution of other shared off-target sites of CNO and C21 such as the 5-HT_2A_ receptors, which are thought to mediate the locomotor suppression after high doses of clozapine (McOmish et al. 2012), should also be taken into consideration.

### Previous indications for sleep-modulatory effects of CNO

Many chemogenetic sleep studies provide adequate control data, which excludes relevant effects of CNO on sleep in the given experimental paradigms (Erickson et al. 2019; Mondino et al. 2021; Takata et al. 2018; Yu et al. 2019; Anaclet et al. 2015; Venner et al. 2016). Many other studies do not present or analyse data of CNO-injected controls but instead only compare CNO versus saline conditions in DREADD-expressing animals. However, occasionally authors have mentioned that the use of CNO injections of up to 10 mg/kg may have affected the sleep of animals in DREADD-free control groups (Funk et al. 2017) and recent work focussing on REM sleep regulation presented statistically significant effects of CNO doses higher than 5 mg/kg in the supplementary data (Varin, Luppi, and Fort 2018). Another well-controlled study found a slight increase in NREM sleep bout duration of control mice injected with 0.3 mg/kg CNO, while all other analysed parameters were unaffected by this small CNO dose (Liu et al. 2021). This finding supports our effect size analysis indicating that NREM sleep bout duration is the sleep architectural parameter most strongly affected by CNO and C21 and that CNO can cause sleep-modulating effects at doses of 1 mg/kg CNO, and possibly below.

### Implications for future use of chemogenetics in sleep research

Previous work has highlighted that several factors such as age, sex and strain of the experimental animals (Manvich et al. 2018), as well as differences in the activity of the cytochrome P450 enzymes converting CNO to clozapine (Mahler and Aston-Jones 2018), might contribute to the variability of side effects of DREADD actuators. In addition, the galenic formulation and use of solvents can alter the pharmacokinetic properties of DREADD actuators (Campbell and Marchant 2018). The hydrochloride salt preparations of CNO used here and in other recent sleep studies (Fernandez et al. 2018; Stucynski et al. 2021) have reduced back-metabolism to clozapine and an improved water solubility and bioavailability compared to equivalent doses of CNO-DMSO preparations (D. C. Allen et al. 2019). The increased bioavailability and thereby supposedly amplified off-target receptor binding might explain that we find sleep-modulating effects of CNO already at doses where other studies have shown no effects. Considering the difficulty to predict behavioural effects of DREADD actuators in a specific experimental paradigm, our study supports the proposal that it is vital to minimise the injected doses of DREADD actuators and to include CNO-injected non-DREADD-expressing control animals in each individual experiment (MacLaren et al. 2016; Mahler and Aston-Jones 2018; Campbell and Marchant 2018). More importantly, our work shows that avoiding clozapine back-metabolism of CNO by using novel DREADD actuators such as C21 does not prevent behavioural side effects. Interestingly, another next generation DREADD agonist, perlapine, which is structurally similar to C21 (Chen et al. 2015; Thompson et al. 2018), is a REM-suppressing sedative and muscle relaxant used in the treatment of insomnia (Ando et al. 1970; S. R. Allen and Oswald 1973; Stille et al. 1973). Chemogenetic approaches that do not require actuators with high chemical similarity to clozapine might provide an alternative for sleep research (Magnus et al. 2019). Considering the complexity of neurotransmitter systems regulating sleep (Saper and Fuller 2017) and the sensitivity of sleep architecture to pharmacological intervention (Riemann and Nissen 2012), testing the effects of new chemogenetic actuators, such as deschloroclozapine (Nagai et al. 2020) or JHU37152 and JHU37160 (Bonaventura et al. 2019), on sleep might prove not only useful for sleep research but could become a valuable tool in evaluating the biological inertness of newly developed chemogenetic actuators.

### Limitations

We would like to highlight that this study was initiated as a control experiment and was not designed to assess dose-dependent effects of CNO. Instead of counterbalancing all experimental conditions, we counterbalanced the saline and the 5 mg/kg CNO condition for the first two sessions to enable a direct comparison between saline and a medium dose of CNO avoiding potential adaptation effects resulting from repeated CNO injections. The decision to conduct a full study and to include C21 injections as an additional condition was only made once visual inspection of pilot data from four animals had indicated relevant effects of CNO on sleep. Because injections of 1 and 10 mg/kg CNO and 3 mg/kg C21 were always performed after the initial two injections, sequence effects for those conditions cannot be excluded. This should particularly be considered in the interpretation of the results on NREM and REM sleep latency, which do not show a dose-dependency. In order to reduce the number of animals used in laboratory research, most of the animals (13 out of 16) were used for combined experiments including other procedures such as light presentation with light emitting diodes (LEDs) or local intracortical microinfusions. The type of experiments, the duration of the rest interval, and the within-subject design of our study make it unlikely that our findings were confounded by previous experiences but this cannot be fully excluded. It should further be noted that the sample size of C21-injected animals was very small (n=7) and all analyses concerning C21 effects should be considered exploratory. To avoid misinterpretation of one sided null-hypothesis testing we provided effect sizes for all pairwise comparisons (*Supplementary Table 3*).

### Conclusions

In conclusion, our study suggests that the DREADD actuators, CNO and C21, have sleep-modulatory effects, which cannot be explained by back-metabolism to clozapine alone but might result from off-target binding to endogenous receptors. While our results require replication in an optimised experimental design, our findings have important implications for the future application of chemogenetics in sleep research and neuroscience. Our study highlights the need to use non-DREADD-expressing controls, even when novel actuators are used that cannot convert to clozapine. Considering the sensitivity of sleep architectural parameters to CNO and C21 demonstrated here, our work reveals a new opportunity of using simple EEG/EMG sleep screening to assess the pharmacological inertness of novel chemogenetic actuators *in vivo*. We are confident that these experimental refinements of the DREADD approach, plus novel technological improvements, will unleash the full potential of this powerful tool in behavioural neuroscience and help pave the way for its clinical application.

## Materials and Methods

### Animals

Sixteen young adult male C57BL/6J mice (age: 113±6 days, weight: 25.2±0.5 g) were used in this study. All animals were sourced internally from the Biomedical Services at the University of Oxford. As this study was originally designed as a control experiment, we did not intend to implant animals exclusively for this project. In order to reduce the number of animals used in laboratory research, 13 animals were implanted for combined sleep experiments involving procedures for this project and related work. All 16 animals used in this study were implanted with a right frontal EEG screw, a reference EEG screw above the cerebellum, and EMG wires in the neck muscles as described previously (Fisher et al. 2016). In addition to this EEG/EMG configuration, which provided the electrophysiological signals analysed in this manuscript, five of the animals were implanted with a frontal left and bilateral occipital EEG screws as well as with an anchor screw in the midline anterior to the frontal EEG screws, which served as a socket for a detachable light-emitting diode (LED) device (Figure 6a); four animals were implanted with a right occipital EEG screw and a cannula (C315I; PlasticsOne) targeted to layer 5 of the primary somatosensory cortex (Figure 6b); four animals received an additional frontal and occipital EEG screw over the left hemisphere as well as a left cerebellar ground screw and a right occipital 16-channel laminar probe as well as a midline frontal anchor screw for the detachable LED device (Figure 6c); three animals received a left frontal and occipital EEG screw (Figure 6d). All animals had a rest interval of at least three days between the previous experiment and this study.

**Figure 6:**
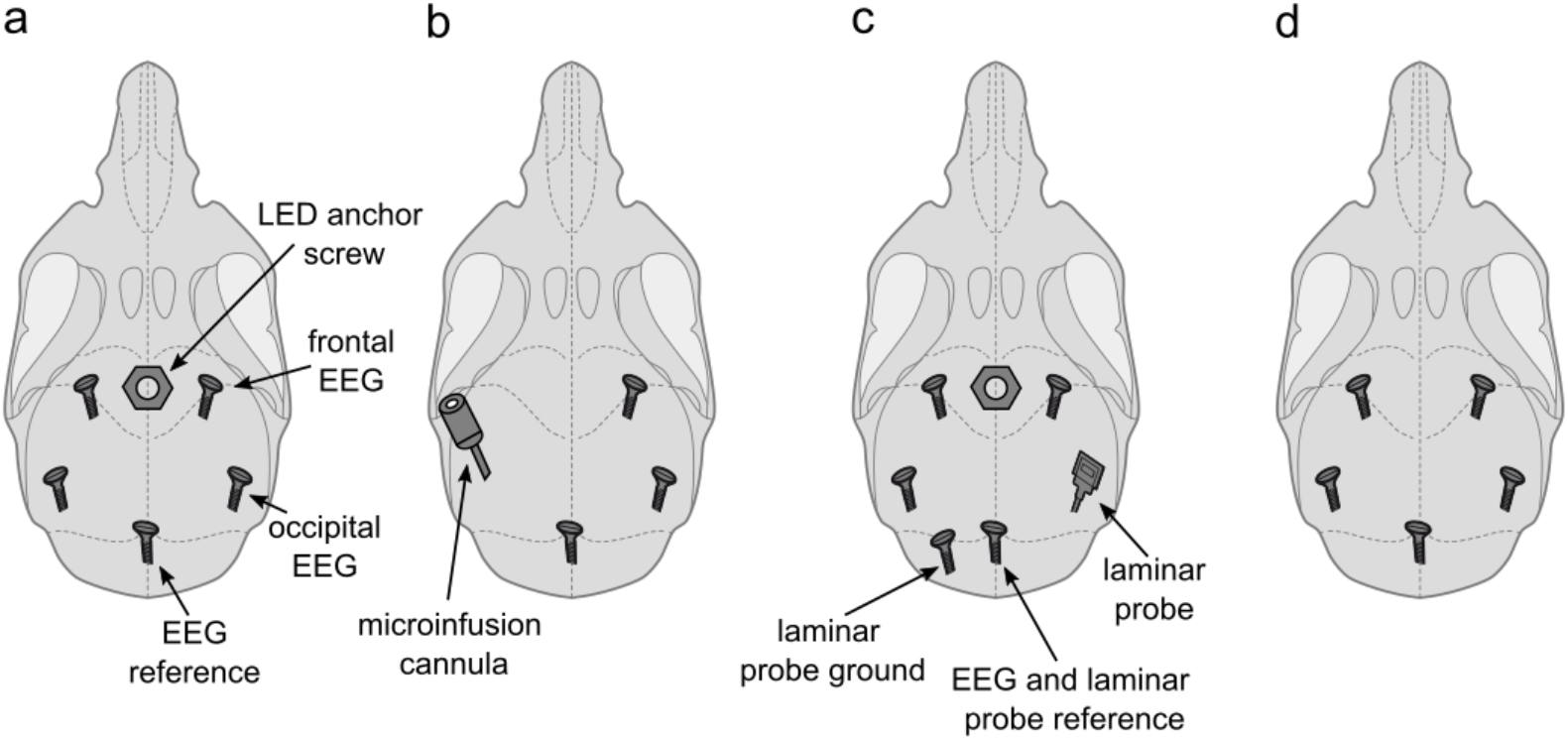
Implant configurations. a) LED anchor screw, bilateral frontal and occipital EEG screws, and cerebellar reference screw. Implant configuration of n=5 mice. b) microinfusion cannula, right frontal and occipital EEG screws, and a cerebellar reference screw. Implant configuration of n=4 mice. c) LED anchor screw, bilateral frontal and left occipital EEG screws, right occipital laminar probe, and cerebellar ground and reference screws. Implant configuration of n=4 mice. d) bilateral frontal and occipital EEG screws, and cerebellar reference screw. Implant configuration of n=3 mice. LED: light-emitting diode. EEG: electroencephalogram.

The nine animals implanted with a socket for the placement of a detachable LED device received flickering light stimulation to one of the eyes combined with a 4-h sleep deprivation on up to four experimental days before being used for our study. All animals had at least three rest days without experimental interventions before the first injections of CNO or saline for the study presented here. The four animals implanted with a cannula in the left primary motor cortex received intracortical microinfusions of bumetanide for a transient and localised blockade of the Na-K-2Cl cotransporter NKCC1 (Kahle and Staley 2008), as well as of VU0436271 for a transient and localised blockade of the chloride potassium symporter KCC2 (Sivakumaran et al. 2015). Two of these animals underwent two intracortical microinfusions and the other two animals underwent three intracortical microinfusions before inclusion in this study. The last infusion took place 12 days before the start of the study presented here. Due to the limit of five injection or infusion procedures on the UK Home Office project license under which our experiments were conducted, these animals could only be subjected to three and two i.p. injections of different CNO doses and saline. The three animals implanted with bilateral frontal and occipital EEG screws were not used for any additional experiments. Only seven animals, the four animals implanted with laminar probes and the three animals used exclusively for this study, were used for C21 injections as this condition was added to the experimental protocol after the pilot data from CNO injections had been obtained.

### Electrophysiological signal acquisition, data processing, and sleep scoring

EEG/EMG recordings were performed using the 128 Channel Neurophysiology Recording System (Tucker-Davis Technologies Inc., Alachua, FL, USA) and the electrophysiological recording software Synapse (Tucker-Davis Technologies Inc., Alachua, FL, USA). Raw data was stored on a local computer in 24-h recording blocks. During the continuous recordings EEG and EMG signals were filtered between 0.1 - 100 Hz, and stored at a sampling rate of 305 Hz. After transfer to an analysis desktop computer, the raw signals were resampled at a sampling rate of 256 Hz using custom-made code in Matlab (The MathWorks Inc, Natick, Massachusetts, USA, version v2020a) and converted into the European Data Format (EDF) as previously described (McKillop et al. 2018). The EDF files were visually scored in individual 4-s epochs by blinded experimenters using the software package Sleep Sign for Animals (SleepSign Kissei Comtec Co., Ltd., Nagano, Japan). If EEG signals contained temporary artefacts due to electrical noise, movements, or chewing, the respective vigilance state was assigned to the respective epoch but the EEG signals were not included in the spectral analysis. Using the Sleep Sign for Animals software, fast Fourier transform routine (Hanning window) with a 0.25 Hz resolution was computed in the frequency range between 0 and 30 Hz for each individual 4-s epoch.

### Clozapine-N-oxide and compound 21 products

To avoid the use of the toxic solvent dimethyl sulfoxide (DMSO), which is typically used for the preparation of CNO products in concentrations of up to 15% (Campbell and Marchant 2018), we opted for the use of a water-soluble salt preparation of CNO, clozapine-N-oxide dihydrochloride (Tocris, Bio-Techne LTD, Abingdon, UK, catalog no.: 6329) dissolved in sterile saline. The dihydrochloride preparation of CNO undergoes less back-metabolism to clozapine but has a higher bioavailability compared to CNO-DMSO as indicated by pharmacokinetic work in rhesus macaques (D. C. Allen et al. 2019). This product has previously been used in sleep studies on mice at concentrations between 1 mg/kg and 5 mg/kg (Fernandez et al. 2018; Stucynski et al. 2021). For C21 injections we used the water-soluble version of DREADD agonist compound 21 (compound 21 dihydrochloride, Tocris, Bio-Techne LTD, Abingdon, UK, catalog no.: HB6124). We chose a dose of 3 mg/kg because a detailed pharmacokinetic assessment of this product at this specific concentration as well as behavioural testing in a 5-choice serial-reaction-time task, which did not reveal any behavioural effects at this dose (Jendryka et al. 2019).

### Experimental design

All 16 animals received a medium dose of CNO (5 mg/kg, 0.25 mg/ml solution) and an equivalent volume of saline in a counterbalanced order with a 72-h rest interval between the first two experimental sessions. Of those 16 mice, fourteen also received a 10 mg/kg CNO injection (0.5 mg/ml solution) and twelve a 1 mg/kg CNO (0.05 mg/ml solution). These two injections were counterbalanced as third and fourth injections. Seven animals also received a 3 mg/kg (0.15 mg/ml) C21 injection in a fifth experimental session. This semi-counterbalanced design was chosen to ensure that at the medium dose condition is counterbalanced with saline injections to circumvent putative habituation or adaptation effects following repeated injection of CNO.

Animals were on a regular light-dark cycle (lights on at 9 am) and intraperitoneal (i.p.) injections were performed within 15 minutes after light onset following a brief health check. The delay between light onset and injection was (mean±s.e.m.): saline: 4.19±0.87 min, 1 mg/kg CNO: 4.58±0.59, 5 mg/kg CNO: 3.52±1.04 min, 10 mg/kg CNO: 4.75±0.59 min, 3 mg/kg C21: 6.01±0.96 min. Individual recording sessions were separated by a rest interval of at least 72 hours. The recording chambers were kept open for approximately 10-15 minutes after the injection to monitor for potential adverse effects. The chambers were then closed and the animals checked remotely at regular intervals for the first 6-12 hours after injection. For data analysis, all recordings were aligned to the time point of injections.

### Sample size determination and power analysis

Sample size and power calculations were performed using G*Power 3.1, an open-source statistical power analysis program (Faul et al. 2007). The sample size was chosen based on previous experiments in our lab investigating the effects of the sedative diazepam on sleep (McKillop et al. 2021), which indicated an effect size of Cohen’s d = 0.90 for the key outcome parameter NREM sleep time. We therefore decided that our study should be sufficiently powered to detect effects of sizes of d=1 designed between saline and individual CNO treatment conditions with a power of 0.9 at the given α-error probability of 0.05. The estimated sample size from this calculation was 12-13 animals per group. Based on experiences from previous EEG studies in mice, we aimed to account for an attrition rate of approximately 20% and decided to include 16 animals in this study.

### Statistical procedures

Data were analysed using MATLAB (version R2020a; The MathWorks Inc, Natick, MA, USA), SAS JMP (version 7.0; SAS Institute Inc. Cary, NC, USA) and IBM SPSS Statistics for Windows (version 25.0; IBM Corp., Armonk, N.Y., USA). Reported averages are mean±s.e.m. For all main effects and interactions an α-error probability of 0.05 was adopted. For the statistical comparison of CNO and saline injections mixed-effect models were calculated using GraphPad Prism (version 9.1.1 for Windows; GraphPad Software, San Diego, California, USA, www.graphpad.com). Significant main effects of treatment conditions or interaction terms were followed up with Dunnett’s adjustment for post-hoc comparisons. In mixed-effect models with multiple time points we analysed acute (0-2 hours post injection) and prolonged (0-6 hours post injection) effects separately. For spectral analysis, EEG/LFP power spectra of individual animals were log-transformed before hypothesis testing. Individual spectral bins were compared between individual CNO treatments and saline used two-tailed t-tests. No correction was applied for post-hoc comparison of spectral data because ANOVAs consisting of 119 EEG spectral bins would be too conservative and reduce statistical power (Achermann and Borbély 1998); thus, α was kept at 0.05 in these cases. Comparisons between the C21 and saline condition were performed using one-tailed *t-*tests to assess whether sleep variables were changed in the same direction as after CNO treatment. For analysis of the percentage of time spent in the three vigilance states, wakefulness and NREM sleep were expressed as the percentage of time in the respective time window, REM was expressed as percentage of total sleep time in the respective time window as in previous work (Huber, Deboer, and Tobler 1999). For sleep architecture analysis, NREM and REM sleep episodes were defined as intervals of at least one minute with allowing an interruption of 4-16 seconds. Brief awakenings are defined as up to 16 second interruptions of sleep by wake-like EEG and EMG patterns (Franken et al. 1991). In all figures, significance levels of post-hoc comparisons are indicated with black asterisks: ‘*’ for 0.05 ≥ p > 0.01; ‘**’ for 0.01 ≥ p > 0.001; ‘***’ for 0.001 ≥ p. Effect sizes are reported as Cohen’s d calculated using the MATLAB function computeCohen_d (Ruggero G. Bettinardi (2020). computeCohen_d(x1, x2, varargin) (https://www.mathworks.com/matlabcentral/fileexchange/62957-computecohen_d-x1-x2-varargin), MATLAB Central File Exchange. Retrieved October 4, 2020). Data from one animal had to be partially excluded because of a defective EEG headstage. This animal had to be excluded from the analysis of brief awakenings due to EMG artefacts, which made it difficult to identify a sudden increase of muscle tone during sleep. The same animal also had to be excluded from the analysis of the 1 mg/kg and 5 mg/kg CNO treatments because of technical issues. Six animals had to be excluded from spectral analysis due to occasional artefacts in the EEG signals.

### Ethical approval

All experiments were performed in accordance with the United Kingdom Animal Scientific Procedures Act 1986 under personal and project licences granted by the United Kingdom Home Office. Ethical approval was provided by the Ethical Review Panel at the University of Oxford. Animal holding and experimentation were located at the Biomedical Sciences Building (BSB) and the Behavioural Neuroscience Unit (BNU), University of Oxford.

### Data and code availability

We encourage readers to further explore the dataset acquired for this publication. Raw data, custom written MATLAB code, and data tables for all figures will be made available on Figshare upon acceptance of the manuscript.

## Supplementary materials

**Supplementary Figure 1:**
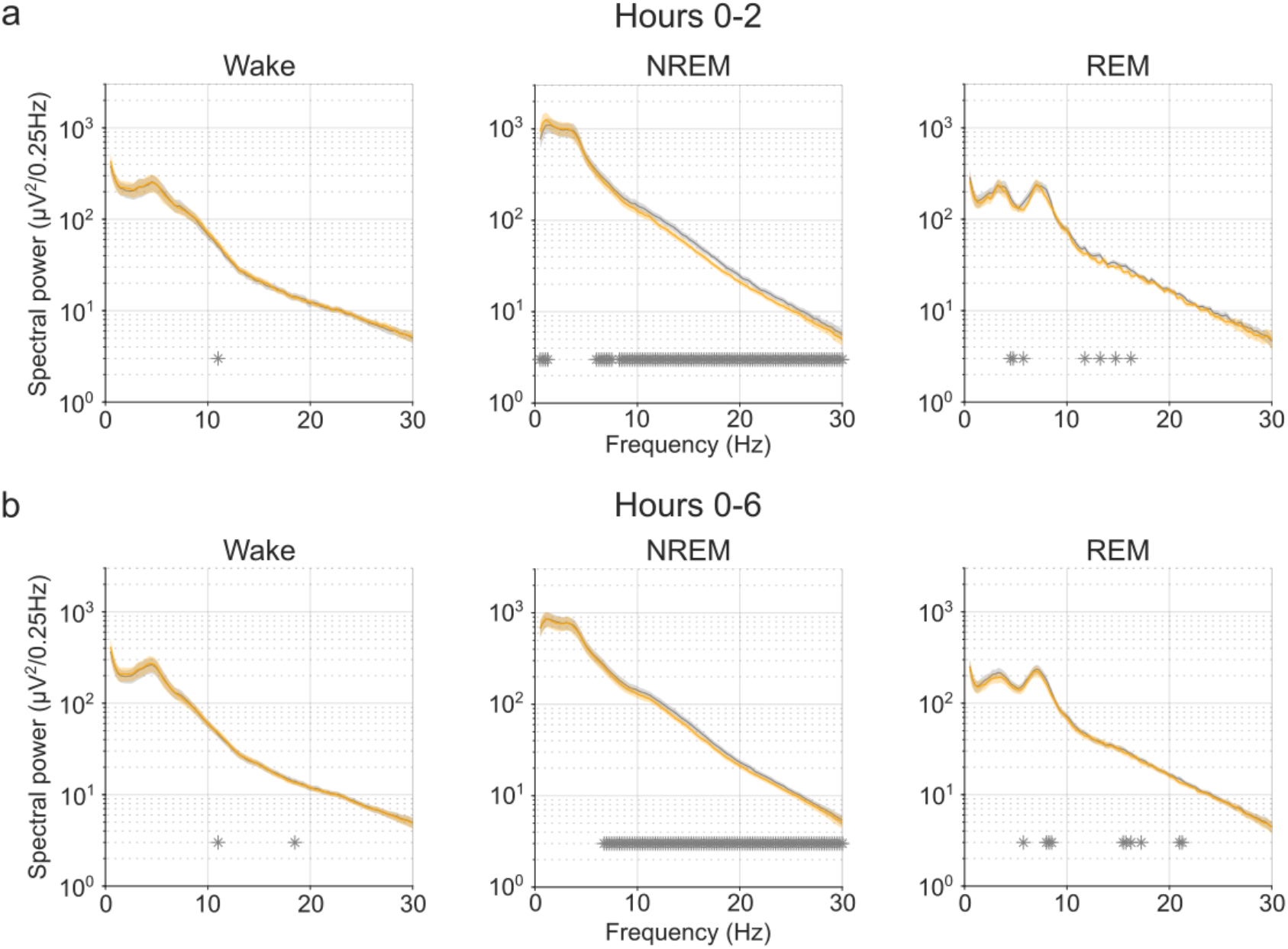
EEG power spectra of wakefulness, NREM, and REM sleep following injections of saline and 5 mg/kg CNO. Frontal EEG spectra in 0.25 Hz bins between 0.5 and 30 Hz arranged by vigilance state and time window. Frequency bins with significant differences in uncorrected post-hoc testing are indicated with grey asterisks. Note the systematic increase of slow frequencies (0-1.25 Hz) in the first 2 hours and the suppression of spectral power in higher frequencies (6.25-30 Hz) in the NREM sleep spectrogram both in the first 2 hours and over the entire observation period. n=10 for saline (grey) and for 5 mg/kg CNO (orange) in spectral analysis. Data are presented as the mean ± s.e.m. (shaded areas). CNO: clozapine-N-oxide. EEG: electroencephalogram. NREM: non-rapid eye movement sleep. REM: rapid eye movement sleep.

**Supplementary Table 1:** Time spent in wakefulness, NREM, and REM sleep after CNO and saline injections.

| | | Treatment condition<br>(mean values ± standard error) | | | | Mixed-effect<br>analysis | | Effect size for post-hoc<br>comparisons (Cohen's d) | | |
| --- | --- | --- | --- | --- | --- | --- | --- | --- | --- | --- |
| Vigilance<br>state | Parameter | Saline<br>(n=16) | CNO<br>1 mg/kg<br>(n=11) | CNO<br>5 mg/kg<br>(n=15) | CNO<br>10 mg/kg<br>(n=14) | F | p | low dose | medium<br>dose | high<br>dose |
| **Wake** |  |  |  |  |  |  |  |  |  |  |
| | 2-h time<br>window (%) | 38.7813<br>±3.2743 | 38.3636<br>±5.7692 | 33.5444<br>±3.8399 | 29.6429<br>±3.9910 | F (1.939,<br>23.91) =<br>1.468 | 0.2504 | -0.0234 | -0.3393 | -0.4815 |
| | 6-h time<br>window (%) | 28.3530<br>±1.3114 | 29.3232<br>±2.5037 | 26.9321<br>±1.5278 | 26.2553<br>±1.7923 | F (1.730,<br>21.34) =<br>0.6888 | 0.4928 | 0.1467 | -0.1715 | -0.2633 |
| **NREM** |  |  |  |  |  |  |  |  |  |  |
| | 2-h time<br>window (%) | 51.9896<br>±2.6391 | 53.7020<br>±4.6576 | 58.7741<br>±3.2742 | 63.7103<br>±3.7117 | F (1.911,<br>23.57) =<br>3.054 | 0.0682 | 0.1157 | 0.4941 | 0.6961 |
| | 6-h time<br>window (%) | 59.0799<br>±1.1641 | 58.9074<br>±1.9870 | 62.1914<br>±1.2667 | 62.6111<br>±1.6784 | F (1.531,<br>18.88) =<br>2.028 | 0.1661 | -0.0361 | 0.4130 | 0.4630 |
| **REM** |  |  |  |  |  |  |  |  |  |  |
| | 2-h time<br>window<br>(% of TST) | 10.4160<br>±0.5877 | 8.1286<br>±1.0534 | 8.4071<br>±0.8351 | 6.5371<br>±0.5937 | F (1.783,<br>21.99) =<br>8.951 | 0.0019 | -0.5802 | -0.5594 | -1.2743 |
| | 6-h time<br>window<br>(% of TST) | 13.1740<br>±0.3774 | 12.3119<br>±0.4884 | 11.1629<br>±0.5722 | 11.0742<br>±0.4606 | F (1.839,<br>22.68) =<br>7.525 | 0.0038 | -0.4697 | -0.7724 | -0.9333 |

**Supplementary Table 2:** Sleep architecture after CNO and saline injections.

|  |  | Treatment condition<br>(mean values ± standard error) |  |  |  | Mixed-effect analysis |  | Effect size for post-hoc comparisons (Cohen's d) |  |  |
| --- | --- | --- | --- | --- | --- | --- | --- | --- | --- | --- |
| Vigilance state | Parameter | Saline (n=16) | CNO 1 mg/kg (n=11) | CNO 5 mg/kg (n=15) | CNO 10 mg/kg (n=14) | F | p | low dose | medium dose | high dose |
| Wake |  |  |  |  |  |  |  |  |  |  |
|  | Longest episode (min) | 31.3292 ±2.3901 | 32.0000 ±3.9528 | 31.6089 ±3.2855 | 31.7952 ±3.5150 | F (2.108, 26.00) = 0.02110 | 0.9825 | 0.0663 | 0.0189 | 0.0298 |
|  | Episode duration average (min) | 11.6562 ±0.9793 | 9.6981 ±0.6636 | 11.0855 ±0.8599 | 11.0939 ±0.9275 | F (2.578, 31.80) = 0.8177 | 0.4777 | -0.4160 | -0.1221 | -0.1355 |
|  | Episode number (n/h) | 1.5313 ±0.1429 | 1.7727 ±0.1107 | 1.4889 ±0.1254 | 1.4286 ±0.0984 | F (2.338, 40.52) = 1.241 | 0.3036 | 0.3046 | -0.0592 | -0.1694 |
| NREM |  |  |  |  |  |  |  |  |  |  |
|  | Longest episode (min) | 16.3458 ±0.6994 | 18.8061 ±1.6012 | 22.1067 ±1.1553 | 26.0905 ±1.5547 | F (2.881, 35.53) = 13.24 | <0.0001 | 0.3562 | 0.9438 | 1.5129 |
|  | Episode duration average (min) | 5.8746 ±0.2822 | 6.1381 ±0.2998 | 7.2894 ±0.3127 | 7.5503 ±0.4268 | F (2.436, 30.04) = 11.64 | <0.0001 | 0.1810 | 1.1430 | 1.1883 |
|  | Episode number (n/h) | 6.3021 ±0.2318 | 5.9697 ±0.3113 | 5.3556 ±0.2444 | 5.2857 ±0.2915 | F (2.476, 30.54) = 7.796 | 0.0010 | -0.2938 | -1.0628 | -1.1989 |
|  | Latency (min) | 30.4958 ±2.9506 | 14.6485 ±2.9015 | 26.1378 ±3.3676 | 25.4762 ±4.6781 | F (2.146, 26.47) = 3.380 | 0.0463 | -1.2021 | -0.3744 | -0.2424 |
| REM |  |  |  |  |  |  |  |  |  |  |
|  | Longest episode (min) | 3.4833 ±0.1631 | 3.2667 ±0.1585 | 3.4622 ±0.1445 | 3.6286 ±0.1480 | F (2.720, 33.55) = 1.101 | 0.3583 | -0.3224 | -0.0272 | 0.1800 |
|  | Episode duration average (min) | 1.9684 ±0.0396 | 1.9699 ±0.0574 | 2.0995 ±0.0662 | 2.2542 ±0.0835 | F (2.330, 40.39) = 4.509 | 0.0132 | 0.0049 | 0.3941 | 1.0034 |
|  | Episode number (n/h) | 2.0521 ±0.0922 | 1.8788 ±0.1441 | 1.8444 ±0.1167 | 1.6786 ±0.0664 | F (2.665, 32.87) = 2.680 | 0.0690 | -0.2975 | -0.3630 | -0.9286 |
|  | Latency (min) | 28.1375 ±2.8814 | 54.8970 ±7.7841 | 36.1778 ±6.0069 | 43.3667 ±4.1811 | F (2.308, 28.46) = 6.985 | 0.0024 | 0.9798 | 0.3517 | 0.8288 |
| Sleep consolidation |  |  |  |  |  |  |  |  |  |  |
|  | NREM before REM onset (min) | 20.9500 ±1.8146 | 31.4970 ±3.5462 | 30.0711 ±3.5065 | 35.9095±3.5029 | F (2.709, 33.41) = 5.071 | 0.0066 | 0.7346 | 0.5683 | 1.0475 |
|  | Brief awakenings (n/h)* | 36.8033 ±3.7206 | 27.6169 ±3.2139 | 20.9897 ±2.7844 | 21.5082±2.0680 | F (1.968, 23.62) = 10.38 | 0.0006 | -0.6271 | -1.0691 | -1.5760 |

\*animal numbers for analysis of brief awakenings: n=15 for saline, n=11 for 1 mg/kg, n=15 for 5 mg/kg, n=13 for 10 mg/kg

**Supplementary Table 3:** Sleep parameters after saline and compound 21 injections.

| Vigilance state | Parameter | Saline (n=7) | C21 3 mg/kg (n=7) | t | p | Effect size (Cohen's d) |
| --- | --- | --- | --- | --- | --- | --- |
| **Wake** |  |  |  |  |  |  |
|  | 2-h time window (%) | 31.4841<br>±4.1581 | 29.3571<br>±3.7073 | t=0.2993,<br>df=6 | 0.3874 | -0.1131 |
|  | 6-h time window (%) | 26.4153<br>±2.1399 | 23.5106<br>±1.9486 | t=0.9230,<br>df=6 | 0.1958 | -0.3489 |
|  | Longest episode (min) | 27.6286<br>±2.5082 | 30.3524<br>±4.1173 | t=0.8184,<br>df=6 | 0.2222 | 0.3093 |
|  | Episode duration average (min) | 10.4205<br>±0.9875 | 10.6269<br>±1.8085 | t=0.1339,<br>df=6 | 0.4489 | 0.0506 |
|  | Episode number (n/h) | 1.5714<br>±0.2463 | 1.4286<br>±0.1776 | t=0.4680,<br>df=6 | 0.3281 | -0.1769 |
| **NREM** |  |  |  |  |  |  |
|  | 2-h time window (%) | 58.9444<br>±2.9443 | 63.2302<br>±3.0788 | t=0.8081,<br>df=6 | 0.2249 | 0.3054 |
|  | 6-h time window (%) | 61.4603<br>±1.8489 | 65.5873<br>±1.6922 | t=1.623,<br>df=6 | 0.0779 | 0.6133 |
|  | Latency (min) | 25.3619<br>±3.2217 | 22.6571<br>±4.8946 | t=0.3844,<br>df=6 | 0.3570 | -0.1453 |
|  | Longest episode (min) | 15.9143<br>±0.7453 | 21.5429<br>±1.6727 | t=2.551,<br>df=6 | 0.0217 | 0.9641 |
|  | Episode duration average (min) | 6.0655<br>±0.5040 | 8.0568<br>±0.5341 | t=2.462,<br>df=6 | 0.0245 | 0.9307 |
|  | Episode number (n/h) | 6.3571<br>±0.2708 | 5.1190<br>±0.3426 | t=2.809,<br>df=6 | 0.0154 | -1.0617 |
| **REM** |  |  |  |  |  |  |
|  | 2-h time window (% of TST) | 9.9182<br>±0.9511 | 7.5927<br>±0.3919 | t=1.829,<br>df=6 | 0.0585 | -0.6915 |
|  | 6-h time window (% of TST) | 12.6278<br>±0.5749 | 10.8374<br>±0.3969 | t=3.234,<br>df=6 | 0.0089 | -1.2223 |
|  | Latency (min) | 26.0095<br>±5.0918 | 37.8952<br>±5.8049 | t=1.474,<br>df=6 | 0.0954 | 0.3718 |
|  | Longest episode (min) | 3.6952<br>±0.1539 | 3.6476<br>±0.2573 | t=0.2047,<br>df=6 | 0.4223 | -0.0774 |
|  | Episode duration average (min) | 2.0054<br>±0.0678 | 2.2206<br>±0.1093 | t=2.229,<br>df=6 | 0.0337 | 0.8425 |
|  | Episode number (n/h) | 2.0000<br>±0.1744 | 1.8333<br>±0.0962 | t=1.000,<br>df=6 | 0.1780 | -0.3780 |
| **Sleep consoli-**<br>**dation** |  |  |  |  |  |  |
|  | NREM before REM onset (min) | 21.6762<br>±3.8776 | 31.5714<br>±3.9007 | t=2.252,<br>df=6 | 0.0326 | 0.8511 |
|  | Brief awakenings (n/h)* | 31.5102<br>±2.6435 | 20.1852<br>±2.6435 | t=2.164,<br>df=5 | 0.0414 | -0.8836 |

\*animal numbers for analysis of brief awakenings: n=6

## Acknowledgements

We thank Prof. Dr. Dennis Kätzel from the University of Ulm, Germany, for his advice on the study design and the preparation of the DREADD actuators; Prof. Dr. Antoine Adamantidis from the University of Bern, Switzerland, for his comments on the manuscript; all members of the laboratory of the Vyazovskiy lab for kind help with surgery assistance and animal care. This work was supported by a travel scholarship from the German Academic Exchange Service (DAAD) and a studentship from the Studienstiftung des Deutschen Volkes awarded to JT as well as a Wellcome Trust PhD studentship 203971/Z/16/Z to LBK. LBK was also supported by a Mann Senior Scholarship in medical sciences by Hertford College, Oxford. LEM was supported by a Novo Nordisk Postdoctoral Fellowship run in partnership with the University of Oxford. LEM was also supported by a Sir Paul Nurse Junior Research Fellowship at Linacre College, Oxford. The laboratory of ZM received funding from the UK Medical Research Council (G00900901), the Royal Society, St John’s College Research Centre, the Anatomical Society and Einstein Stiftung. ZM is an Einstein Visiting Fellow at Charité-Universitätsmedizin Berlin (host B. Eickholt for 2020–2024), and lead researcher at Oxford Martin School, University of Oxford. This work was further supported by a Wellcome Trust Strategic Award (098461/Z/12/Z), a John Fell OUP Research Fund grant (131/032) and Medical Research Council (UK) grants MR/N026039/1 and MR/ S01134X/1.

## Author contributions

JT, AHS, ZM, CJA, VVV, and LBK designed the study. JPM, EM, LEM, HA, and LBK performed surgeries. JT, JPM, EM, LEM, LBK conducted the experiments. EM, AHS, and LBK prepared the drug solutions. JT, JPM, EM, LEM performed the sleep scoring. JT, LEM, SMS, VVV, and LBK analysed the data. JT, ZM, CJA, VVV, and LBK wrote the manuscript with input from all authors. All authors discussed the results and commented on the manuscript.

